# Targeting human M2 macrophages with antibodies in optimized 3D tumor spheroids

**DOI:** 10.1101/2025.09.16.676563

**Authors:** Chloé Bazile, Fabien Gava, Flavien Raynal, Marcin Domagala, Julie Bordenave, Laetitia Ligat, Manon Farcé, Corinne Bousquet, Vera Pancaldi, Véronique Gigoux, Mary Poupot

## Abstract

Tumor microenvironment (TME) constituents, including tumor-associated macrophages (TAMs), are now well known to have a significant impact on tumor development. TAMs can be predominant cells in the TME, able to promote cancer cell proliferation, resistance to treatments and immunosuppression. Inactivating these TAMs in tumors by their depletion or their repolarization into a specific anti-tumor, inflammatory phenotype constitutes a significant challenge in immuno-oncology. Several tools and methods to target TAMs have been proposed but they often show low specificity for pro-tumoral macrophages and target myeloid cells too broadly. This research area is therefore in full expansion and warrants the development of appropriate study models. Given the current global effort to reduce the use of *in vivo* approaches, developing *in vitro* models that mimic the behavior of TAMs in the TME has become a priority. In this study, we focused on targeting pro-tumoral TAMs in a simple *in vitro* model recapitulating cancer cell proliferation and TAMs specific phenotype. We developed a 3D model consisting of cancer cells and pro-tumoral macrophages, in which these macrophages promote tumor cell proliferation, maintaining an immunosuppressive phenotype. We showed that, in this model, macrophages can be easily targeted and killed with specific antibodies even in the central regions of the spheroid. This model could be a first *in vitro* approach to screen new TAM depletion or depolarization tools before application in *in vivo* models.

## INTRODUCTION

Cancer remains incurable unless complete surgical removal is achieved, largely due to tumor resistance mechanisms rooted in the tumor microenvironment (TME). Among the TME components, tumor-associated macrophages (TAMs) play a crucial pro-tumoral role by promoting cancer cell survival, proliferation, migration, and protection from immune responses and therapies [1]. TAMs can constitute a significant portion of the tumor mass—up to 30-50% in lung cancer—and are strongly linked to tumor progression and aggressiveness [2].

Macrophages within tumors exist on a spectrum between two main phenotypes: M1 and M2 [3]. Early-stage tumors often have M1 macrophages, which are anti-tumoral and able to kill cancer cells. As the tumor evolves, the microenvironment shifts macrophages toward the M2 phenotype, which supports tumor growth and suppresses immune responses. These M2-like TAMs contribute to poor clinical outcomes and resistance to chemotherapy and immunotherapy [4].

Given their pivotal role in tumor progression and treatment resistance, TAMs have become a key therapeutic target. Some strategies focus on preventing TAMs recruitment, reprogramming their phenotype from M2 to M1, blocking their tumor-supporting functions, or depleting them entirely [5,6]. These approaches include the use of cytokines, small molecules, nanoparticles, and immune-modulating antibodies. However, issues remain in delivering these agents effectively within tumors and understanding their impact on cancer cells.

To address these challenges, reliable *in vitro* models are critical before *in vivo* testing. Several 3D tumor models incorporating TAMs have been developed to better replicate the TME complexity. For example, multicellular spheroids combining cancer cells with mesenchymal stem cells and macrophages have been used in osteosarcoma studies [7]. Many models use the THP-1 cell line as a macrophage surrogate, activated non-specifically, to simulate TAMs (PDAC: [8]; NSCLC: [9,10]; Carcinoma: [11]). Although convenient, THP-1 macrophages do not fully mimic the physiological state of TAMs.

Other approaches involve adding monocytes to tumor-fibroblast spheroids, to promote their differentiation into M2 macrophages, though the phenotype varies with cancer type and conditions. Use of conditioned media to promote M2 markers shows limited success, as key markers like CD206 may not be consistently induced [12]. More complex models incorporate pancreatic stellate cell-derived fibroblasts and endothelial cells, demonstrating that IL-10 and endothelial signals are essential for M2 polarization, marked by CD163, CD206, PD-L1, and CD40 expression [13].

Some studies used patient-derived cells, mixing tumor explants with autologous macrophages, which showed TAM infiltration and maintenance of immunosuppressive phenotypes within spheroids [14]. These models closely resemble human tumors, while being limited by the availability of matched patient samples. Similarly, co-cultures of head and neck cancer cells with monocytes from peripheral blood mononuclear cells (PBMCs) successfully induced M2 macrophage differentiation expressing CD206 and IL-10 [15].

Murine models have also been considered, for example using bone marrow-derived macrophages (BMDMs) co-cultured with breast cancer cells to test macrophage reprogramming agents such as anti-CSF1R antibodies or IFN-γ and Pam3SCK4, demonstrating antibody penetration and macrophage phenotype modulation inside spheroids [16,17]. However, species differences limit direct translation to human tumors.

In this study, we developed a simple 3D co-culture model to study macrophage behavior and target TAMs in a lung cancer-like environment by producing spheroids with A549 lung cancer cells and monocytes or polarized macrophages. Various cell ratios and matrix types were tested to optimize the model. We evaluated cell viability, proliferation, macrophage phenotype, and spatial distribution. Using high-resolution microscopy, we demonstrated that antibodies could penetrate the spheroids to specifically target macrophages, supporting therapeutic studies.

## RESULTS

### Monocytes do not polarize toward M2 macrophages in 3D co-cultures with cancer cells

To develop a robust 3D lung cancer model for studying macrophage behavior and targeting TAMs, we first optimized the cellular density for spheroid formation using A549 lung cancer cells. Spheroids formed with 1,000 to 10,000 cells in ultra-low-attachment U-shaped bottom plates showed growth up to the seeding density of 2,500 cells, as indicated by the increased number of cells and spheroids sizes (Supplementary Figure S1). At higher densities, the proliferation was slowed down with no impact on the area.

According to those results, 2500 cells seeding condition was retained for further experiments. We generated 3D spheroids combining 2,500 A549 cancer cells with 2,500 primary myeloid cells (ratio 1:1) - either monocytes or M2-polarized macrophages (M2M) derived from healthy donor monocytes. The spheroid’s growth was analyzed, as previously, by counting cancer cells for 7 days and measuring the spheroid’s area (Figure 1A-C). Over 7 days, cancer cell viability, defined by cell counting and the percentage of viable cells analyzed by flow cytometry, and spheroid’s area increased significantly in the presence of M2M compared to monoculture or co-culture with monocytes (Figure 1A-C), indicating the pro-tumoral role of M2M in promoting cancer cell proliferation within spheroids.

**Figure 1:**
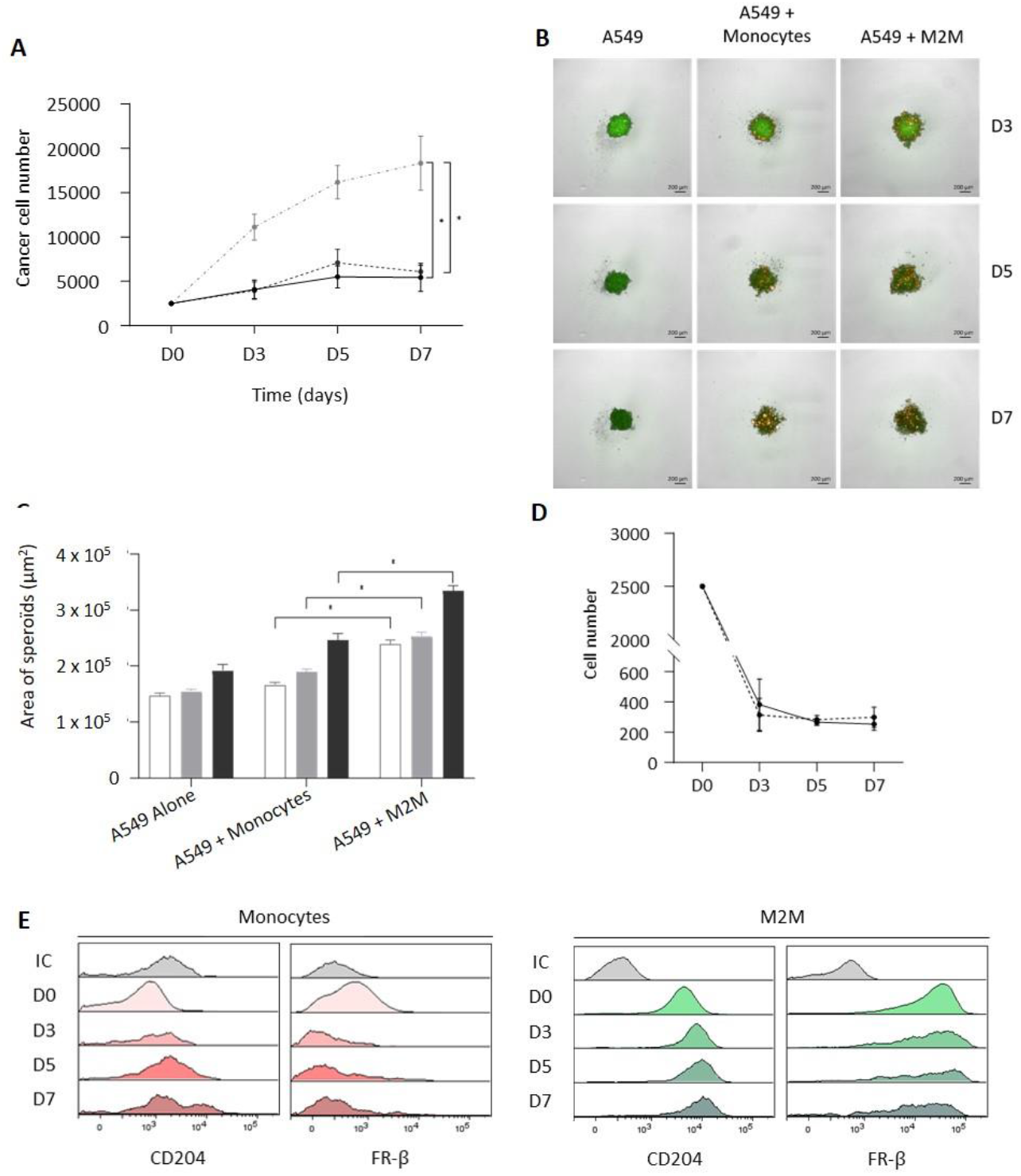
Monocytes in 3D co-culture with tumour cells do not acquire an M2 phenotype. **A-B** - Cancer cells’ (A549 cell line) proliferation in 3D co-cultures with monocytes or M2 derived macrophages (M2M) for seven days: **A-** Number of A549-CFSE^+^ counted on a Malassez chamber (black: A549 alone (n=4), black dotted line: A549 + monocytes (n=4), grey dotted: line A549 + M2M (n=12)). Results are expressed as the mean ± standard error of the mean (sem). Statistical analysis was performed with a two-way anova. **B-** Representative images of a projection of the 3D co-cultures by 2D imaging (Operetta) at different days for the three conditions (A549 in green and monocytes or M2M in orange). **C-** Spheroid area (white: day 0, grey: day 5, black: day 7 (n=30)), expressed as the mean ± sem. Statistical analysis was performed using the Wilcoxon test. **D** - Number of M2M-CMTMR^+^ counted on a Malassez chamber (black dotted line) or monocytes-CMTMR^+^ (black solid line), expressed as the mean ± sem, in the 3D co-cultures with A459 for seven days. **E** - Flow cytometry analysis of CD204 and FR-β expression by M2M or monocytes in 3D co-cultures with A549 for seven days (one representative experiment, isotypic control is shown in grey).

All together, these results indicate that M2M presents pro-tumoral activity promoting the proliferation of cancer cells in spheroids. Of note, the number of viable myeloid cells decreased in spheroids from day 0 to day 3, both for M2M or monocytes, but stabilized afterward (Figure 1D). Importantly, monocytes did not polarize into M2 macrophages during co-culture, as shown by the lack of M2 markers CD204 and FR-β even after 7 days (Figure 1E). This absence of pro-tumoral polarization aligns with the lack of monocyte-induced enhancement in cancer cell proliferation. These findings demonstrate that while M2M supports tumor growth, monocytes alone do not acquire pro-tumoral properties in this model.

### Impact of the matrix on cancer cell proliferation in the 3D co-cultures with M2 macrophages

Next, the effect of matrix support was assessed in co-cultures with M2M. 2,500 A549 cells, pre-labeled with CTV (cell tracer of viability), were mixed with 2,500 M2M to form spheroids either in the absence or in the presence of Matrigel at three different concentrations. The decrease in CTV labeling over seven days indicates that the cancer cells proliferated under all conditions (Supplementary Figure S2). The number of cancer cells determined by the counting of total cells in the dissociated spheroids and the percentage of viable cancer cells using flow cytometry, showed no significant effect of Matrigel on cancer cell proliferation from day 1 to day 7 (Figure 2A). However, cancer cell viability was slightly reduced in the presence of high Matrigel concentrations (Figure 2B), and microscope images showed that spheroids were less structured when cultured with high Matrigel concentrations (Figure 2C). In addition, we observed that the number of M2M tended to decrease more markedly in spheroids grown with Matrigel compared to those grown without Matrigel (Figure 2D). Taken together, these results indicate that Matrigel did not promote spheroid growth. Therefore, A549/M2M spheroids were cultured in the absence of Matrigel in subsequent experiments.

**Figure 2:**
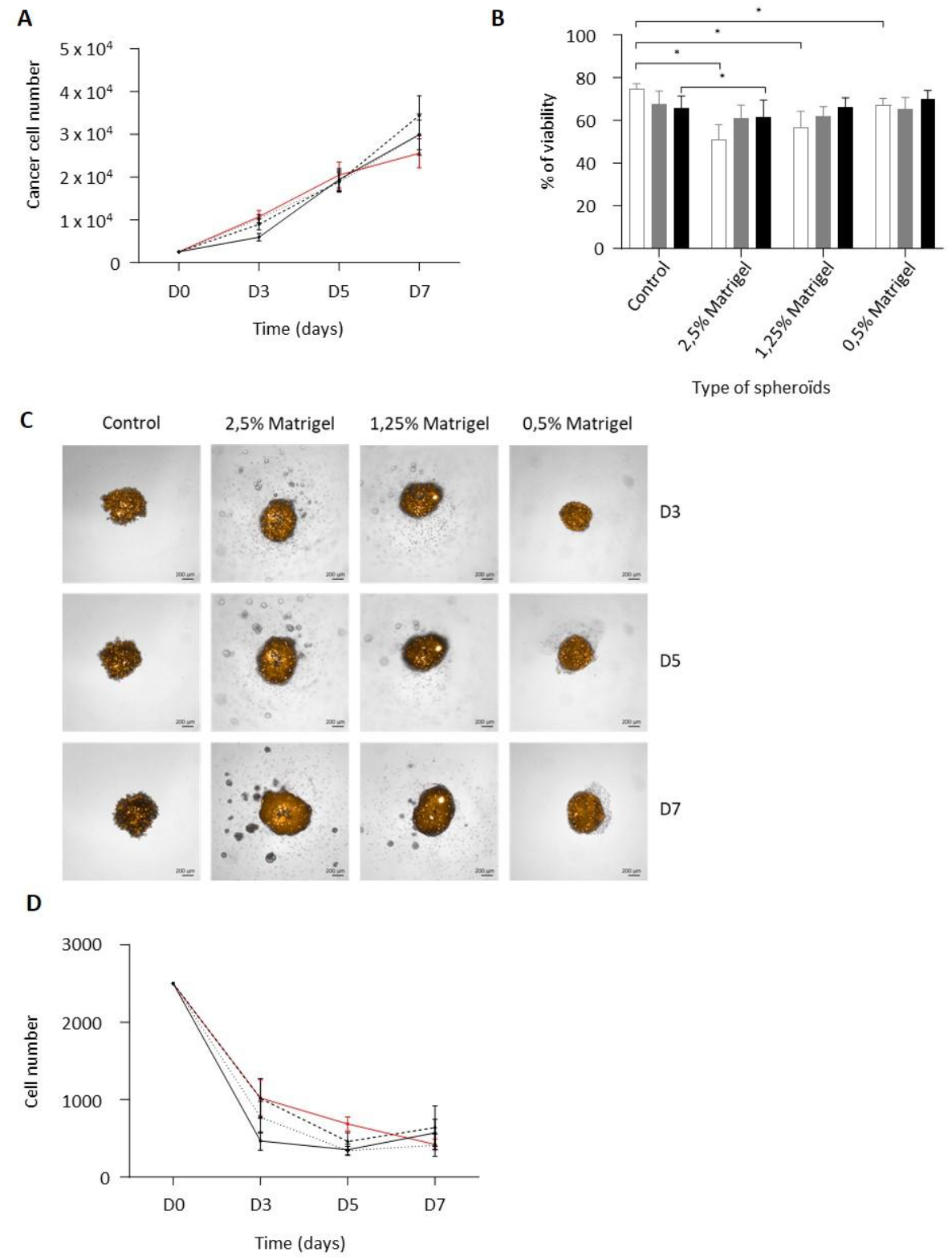
Impact of the matrix support on the 3D models. **A-** Number of A549-CTV^+^ counted on a Malassez chamber, expressed as the mean ± sem, in A549/M2M spheroids for 7 days (**A:** red line: Control ; black solid line: 2.5% of Matrigel ; black dashed line: 1.25% of Matrigel; black dotted line: 0.25% of Matrigel). **B**-A549 viability percentage in A549/M2M spheroids analyzed using Annexin-V/7-AAD labeling by flow cytometry (**B**: white: day 0, grey: day 5, black: day 7). Results are expressed as the mean ± sem. Statistical analysis was performed using paired t-tests. **C-D** - 3D co-cultures of A549 with M2M for 7 day (6 independent experiments): Representative images of a projection of the 3D co-cultures by 2D imaging (Operetta) at 5X objective (**C**: A549 in black and M2M in orange) and quantification of M2M number (**D**: red line: Control ; black solid line: 2.5% of Matrigel ; black dashed line: 1.25% of Matrigel; black dotted line: 0.25% of Matrigel).

### The M2 phenotype is maintained in 3D model with cancer cells

We then investigated whether co-culturing M2 macrophages (M2M) with A549 lung cancer cells influences the M2 phenotype. We compared the viability and phenotype of M2M cultured alone or with cancer cells under 2D and 3D conditions for seven days. M2M viability remained similar across all culture conditions, including monocultures and co-cultures in both 2D and 3D (Supplementary Figure S3). We then analyzed the expression of M2 markers (CD206, CD204, CD209, FR-β, CD163) and M1 markers (CD86, CD64) by flow cytometry after seven days of spheroid growth (Figure 3A–B). CD163, CD206, and CD209 levels were stable across all conditions, while CD204 expression increased in both 2D and 3D co-cultures compared to monoculture. FR-β expression notably increased in 2D co-cultures compared to 3D or monocultures (Figure 3A–B). Expression of M1 markers CD86 and CD64 remained steady in all conditions (Figure 3A, Supplementary Figure S4).

**Figure 3:**
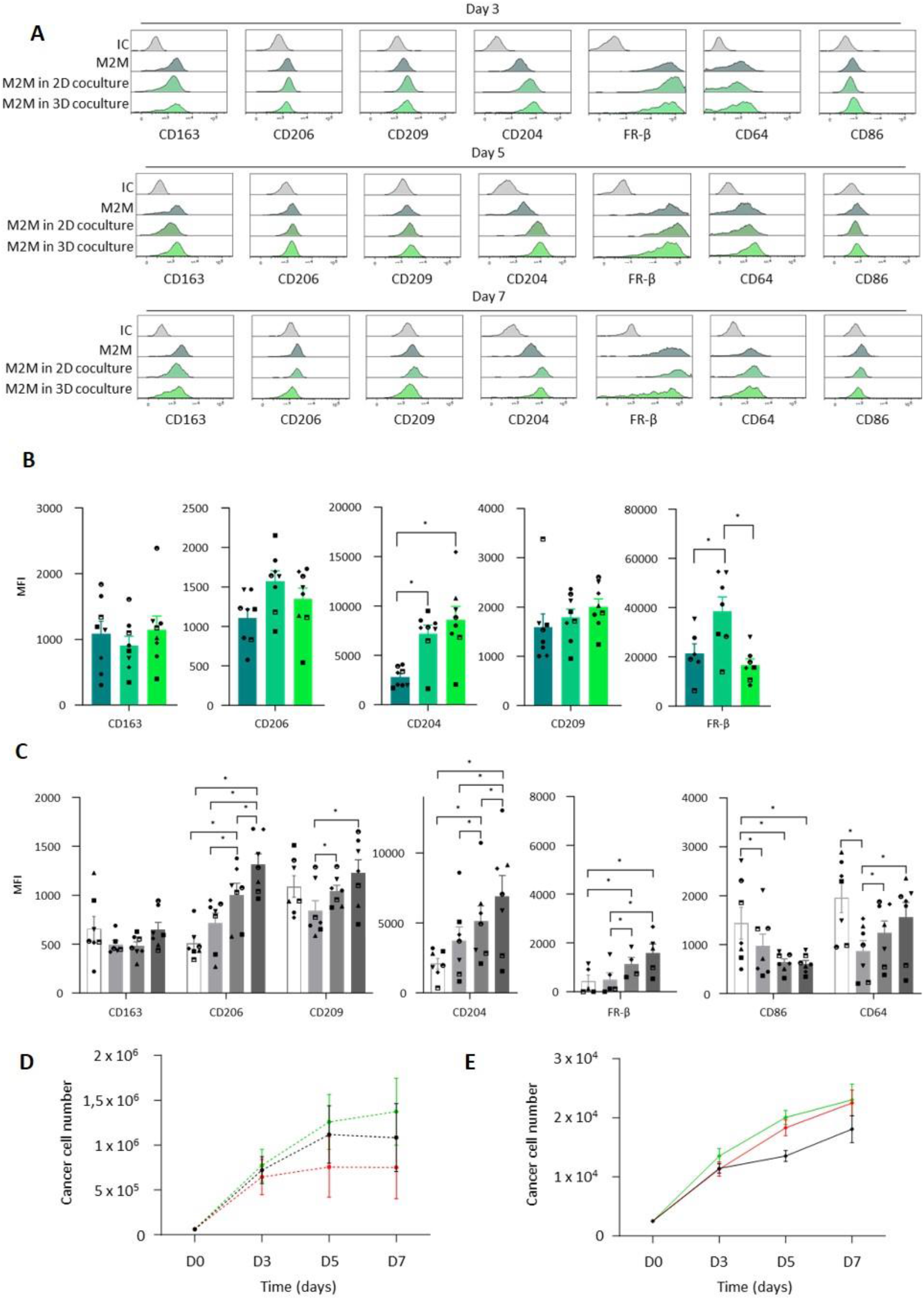
Macrophages phenotypes in the 3D co-culture model. **A-B-** Expression of CD163, CD206, CD209, CD204, FR-β, CD64 and CD86 measured by flow cytometry on M2M (grey: isotypic control; dark green: M2M alone; green: M2M in 2D co-culture; light green: M2M in 3D co-culture). **A-** one representative experiment; **B-** Quantification of CD163, CD206, CD204, CD209 and FR-β expression by flow cytometry at five days of A549/M2M spheroids growth. Results are expressed as the mean ± sem of seven independent experiments. Statistical analysis was performed using Wilcoxon test or paired t-test depending on the distribution of the marker values. **C**: Analysis of M2 markers (CD163, CD206, CD209, CD204, FR-β) and M1 markers (CD86, CD64) at day 0 (white), day 3 (light grey), day 5 (grey) and day 7 (dark grey) in 3D A549/M1M co-culture. Results are expressed as the mean ± sem of seven independent experiments. Statistical analysis was performed using Wilcoxon test or paired t-test depending on the markers. **D-E-** Number of A549 in 2D (**D**) / 3D (**E**) co-cultures for 7 days (black line: A549 alone; green and red line: A549 cultured with M2M or M1M respectively). Results are expressed as the mean ± sem of 8 independent experiments with M2M and 7 independent experiments with M1M.

To validate the model, M1 macrophages (M1M) were co-cultured with A549 cells in 3D for seven days to assess phenotypic changes (Figure 3C). We observed a significant expression rise in CD206, CD204, and FR-β and a decrease in CD86, indicating M1M depolarization toward an M2-like phenotype over time (Figure 3C). We also assessed how M1M and M2M affect A549 proliferation in 2D and 3D co-cultures (Figures 3D–E). In 2D, A549 proliferation seems to decrease in the presence of M1M compared to monoculture, supporting the maintained anti-tumoral function of M1M, despite slight increases in CD204 and FR-β (Supplementary Figure S5). Conversely, in 3D cultures, A549 proliferation was similar with either M1M or M2M, suggesting that M1M lose their anti-tumoral activity and adopt a pro-tumoral, M2-like behavior consistent with their phenotypic changes (Figure 3C). Importantly, M2M tends to increase A549 proliferation in both 2D and 3D cultures, confirming their pro-tumoral activity.

These results demonstrate that M1M and M2M differently influence cancer cell proliferation and reveal distinct behaviors of M1M depending on the culture conditions, while M2M maintain a stable pro-tumoral phenotype. Additionally, A549/M2M 3D spheroids provide a relevant in vitro lung cancer model to study M2M behavior, their impact on tumor cell proliferation, and strategies for targeting M2M.

### Targeting M2 macrophages in 3D *in vitro* models with cancer cells

First, we clearly showed that M2M were distributed throughout the spheroids, including in the center, and that they were labeled by all tested antibodies after 72 hours of incubation (Figure 4A). Moreover, we demonstrated that M2M can be specifically targeted inside living spheroids with different antibodies: CD163, CD206, CSF1-R and FR-β. To optimize M2M targeting, we performed a time-course incubation of living spheroids with fluorescent anti-CD206 and anti-FR-β antibodies for periods ranging from 6 to 72 hours. We observed that M2M targeting by anti-CD206 could be detected after just 6 hours of incubation. However, 72 hours of incubation resulted in a higher number of macrophages stained with both antibodies, as shown by the overlay of red (CMTMR), blue (anti-CD206), and white (anti-FR-β), which appears pink (Figure 4B). We also demonstrated efficient M2M targeting with fluorescent anti-CSF-1R and anti-CD163 antibodies (Supplementary Figure S6). To further confirm the colocalization of CD206, FR-β, and CMTMR staining, colocalization profiles were generated, revealing a clear overlap of the three colors, with no overlap with green, which marks tumor cells (Figure 4C). A similar colocalization profile was observed when staining with CD163 instead of CD206 (Supplementary Figure S6). Finally, to better analyze M2M targeting, we created in-silico 3D models of the labeled spheroids using Imaris software (Figure 4D).

**Figure 4:**
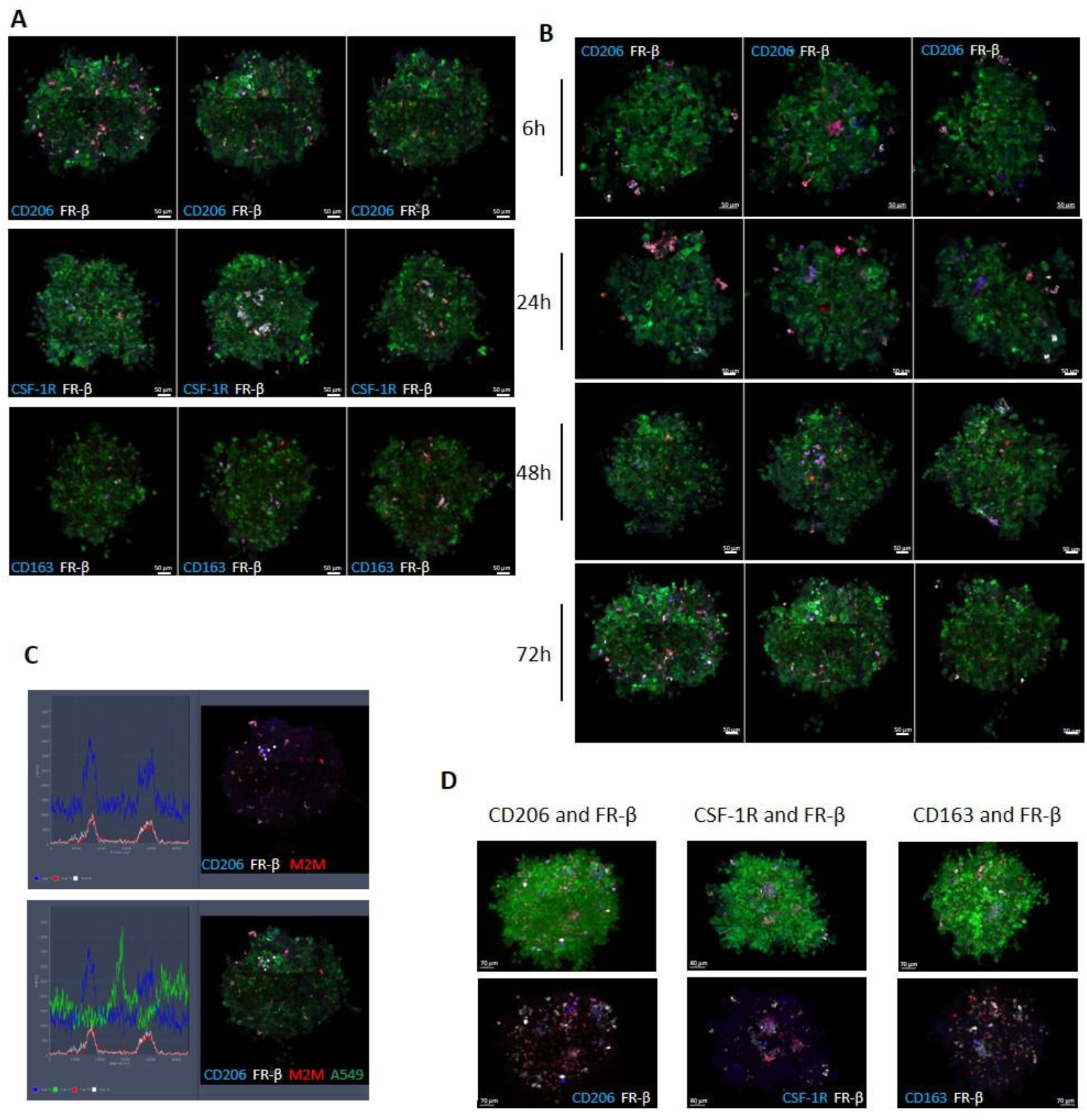
M2-polarized macrophages targeting in 3D models. **A-** Confocal images of 3D models of GFP A549 (green) and M2M-CMTMR (red) incubated with anti-CD206 (blue), anti-CSF-1 (blue), anti-CD163 (blue) and anti-FR-β (white) during 72h at three different z-position inside the spheroid. **B**-Confocal images of 3D models of A549 (green) and M2M (red) incubated with anti-CD206 (blue) and anti-FR-β (white) during different time. **C-** Colocalization profile for 3D model stained with anti-CD206 (blue) and anti-FR-β (white) during 72h. **D**-3D reconstruction of spheroids after 72h staining with different antibodies using Imaris.

### In-silico reconstruction of 3D distribution of M2M in the spheroids

Starting from z-stack confocal images, 3D representations of the entire spheroid can be reconstructed (Supplemental video), making it easier to count cells that are co-labeled with different antibodies (Figure 5A). Using modeling software, we can precisely quantify both the total number of M2M within the spheroid and the subset targeted by specific antibodies. To increase confidence in M2M quantification and compare different analytical approaches, we assessed two distinct in silico segmentation strategies for quantifying antibody-labeled macrophages within spheroids: an automatic procedure and a manual method.

**Figure 5:**
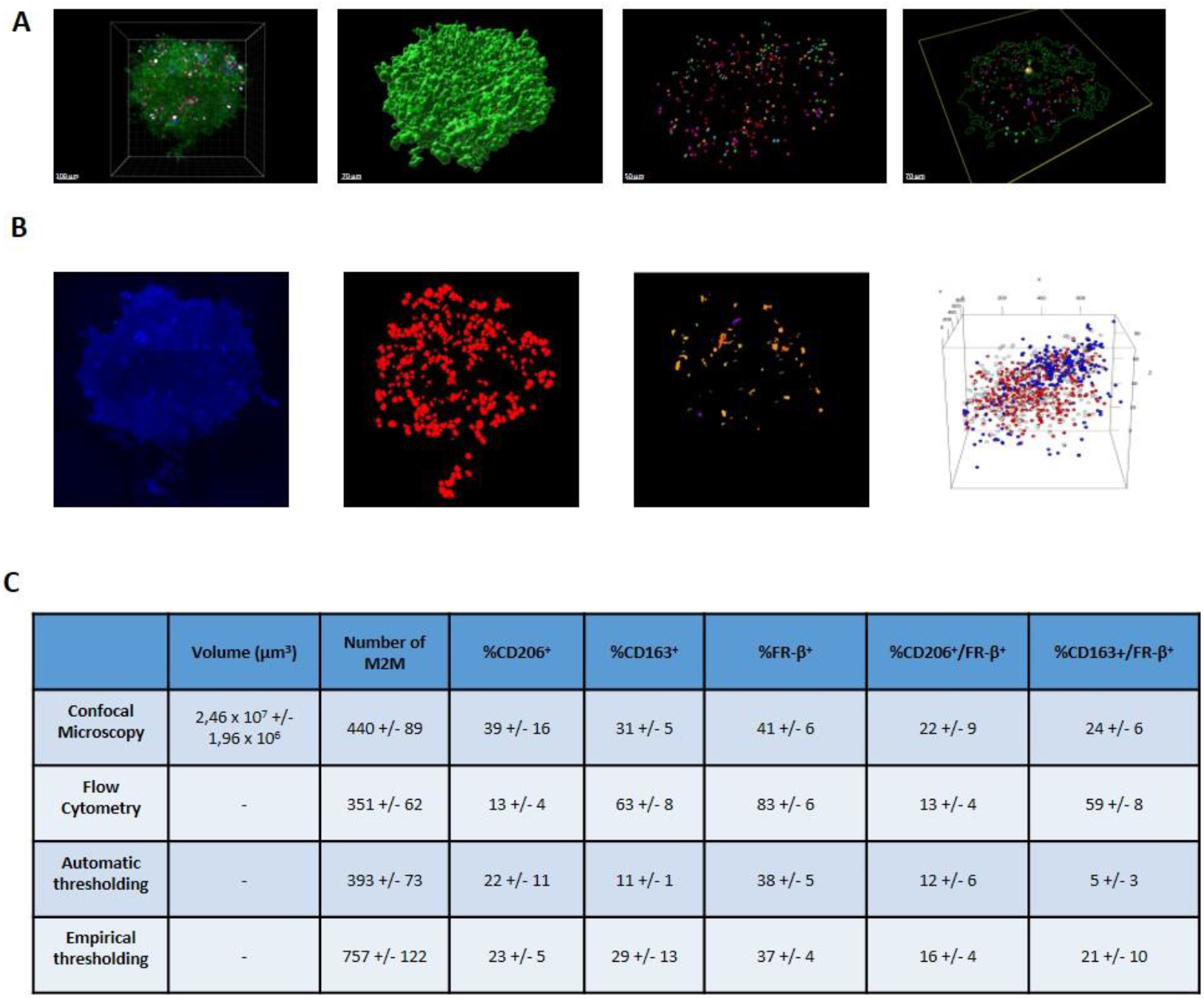
Reconstructions of 3D models of co-culture spheroids from z-stack confocal images. **A-** Imaris reconstruction of 3D models of co-culture spheroids: with all the fluorescences, green for A549, red for CMTMR^+^ macrophages, blue for CD206 and white for FR-β (1), with spots resulting from the 3D reconstruction corresponding to CMTMR/CD206/FR-β staining for macrophages and green for A549 (2), without taking in account the green A549 (3), with a section of the spheroid (4). **B-** In silico signal segmentation: starting from raw fluorescence images (e.g., CD206) (1), signals are extracted using defined thresholds (2), a 3D segmentation algorithm (plugin 3D object counter, Fiji) identifies signal volumes across z-stacks (3). Red (M2M), blue, and white fluorescent signals are then reconstructed in a shared 3D space (4) to calculate distances and determine antibody labeling for each individual M2M. **C-** Mean of the count +/-sem of M2M by different methods of confocal microscopy (n=5), FACS (n=8), in silico 3D image analysis using automatic thresholding (n=5) and empirical thresholding (n=5). Quantification includes total M2M, M2M stained with CD206 (n=3), CD163 (n=2) and FR-β (n=5), and of M2M with colocalization of mAb staining, CD206 and FR-β or FR-β and CD163.

Using automatic thresholding (see Supplemental for methods), which included gamma correction and default intensity thresholds, the average number of M2M per spheroid was 393 ± 73, with 22 ± 11% of cells positive for CD206, 11 ± 1% for CD163, and 38 ± 5% for FR-β. Double labeling revealed 12 ± 6% of M2M positive for CD206 and FR-β, and 5 ± 3% positive for CD163 and FR-β.

In contrast, an empirical thresholding method—based on manual adjustment of signal intensity and without gamma correction—resulted in a higher estimated number of M2M (757 ± 122). The proportions of antibody-positive cells were 23 ± 5% for CD206, 29 ± 13% for CD163, and 37 ± 4% for FR-β. Double-positive cells were more frequent with this method: 16 ± 4% for CD206^+^/FR-β+ and 21 ± 10% for CD163^+^/FR-β^+^.

These findings reinforce the reliability of M2M quantification with labeling strategies. While both segmentation methods are justifiable, the choice of strategy can influence results due to their respective strengths and limitations. The empirical method tends to be more inclusive, detecting more cells by capturing low-intensity signals, whereas the automatic method applies stricter criteria, resulting in a more consistent but more stringent selection.

### Depletion of M2M with macrophage-killing molecule BLZ945

We also investigated the efficacy of therapeutic targeting of M2M in 3D spheroids using the well-known molecule BLZ945, which targets CSF-1R and induces macrophage death [18]. First, we confirmed the expression of CSF-1R on macrophages within the spheroids (Figure 6A). We then evaluated the lethal concentration of BLZ945 on M2M in 2D monocultures and found that 1 µg/ml of BLZ945 killed approximately 50% of M2M after 96 hours of incubation (Figure 6B). The same experiments were performed on A549/M2M spheroids treated with 1 µg/ml BLZ945 (Figure 6C). FACS analysis showed that 1 µg/ml of BLZ945 eliminated all macrophages in the spheroids by day 4 of treatment, without affecting the viability of cancer cells (Figure 6C). Finally, confocal microscopy confirmed these results by showing a reduction in the number of M2M, consistent with the flow cytometry data (Figure 6D). Actually, the mean number of M2M was decreased from 282 to 84 when spheroids were cultured with BLZ945.

**Figure 6:**
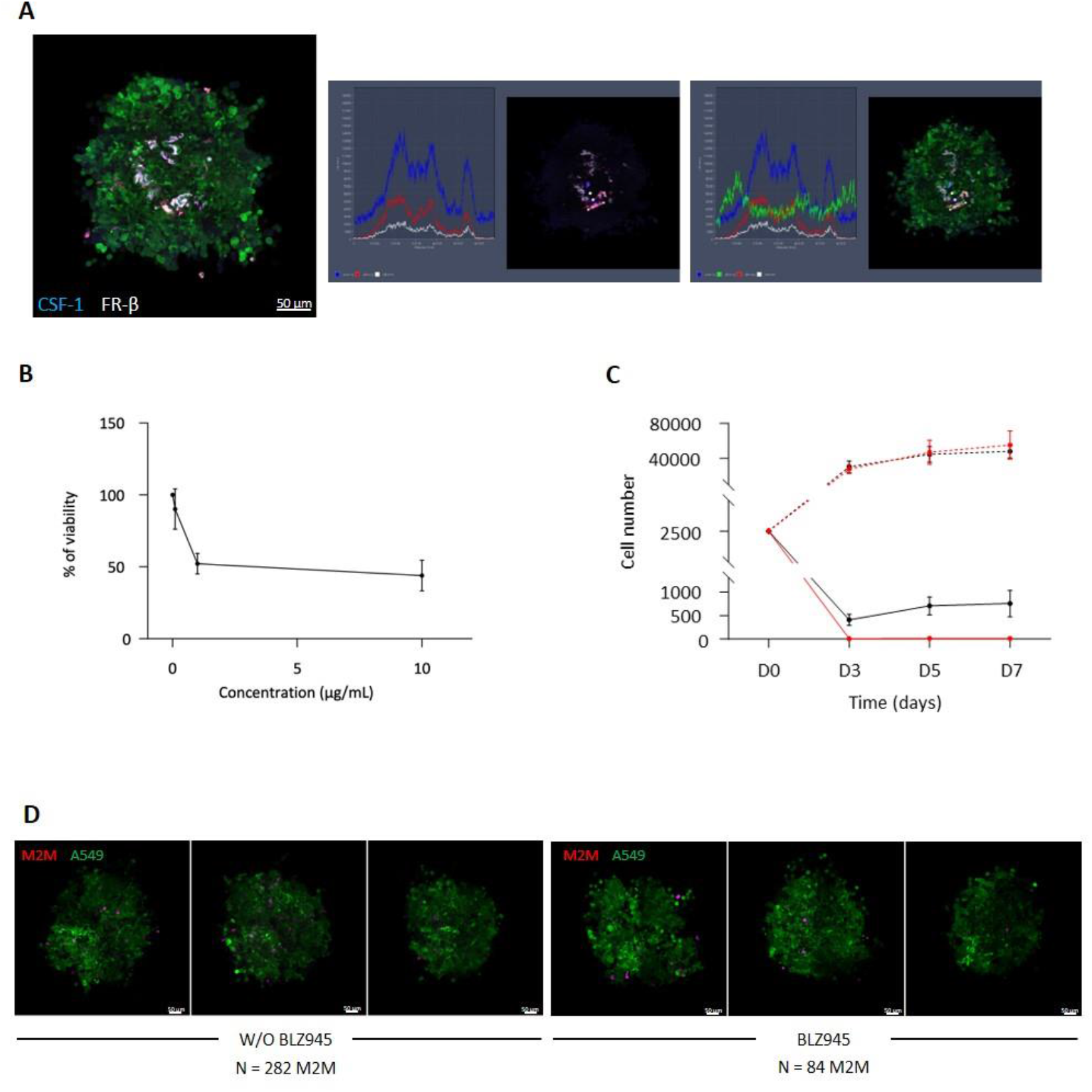
M2 polarised Macrophage depletion in 3D models. **A-** Confocal images of 3D co-culture spheroids of A549-GFP (green) and M2M-CMTMR (red) incubated with anti-CSF-1 (blue) and anti-FR-β (white) during 72h with the colocalization profile. **B-** Toxicity of BLZ945 in 2D monocultures of M2M at different concentrations. **C-** Number of M2M (solid line) or A549 (dotted line) in 3D co-cultures for 10 days post treatment (black line: control; red line: with BLZ945). Results are expressed as the mean ± sem of 3 independent experiments. **D-** Confocal images of 3D models of GFP A549 (green) and M2M-CMTMR (red) incubated with BLZ945 at 1 µg/mL during 4 days or not (six independent experiments).

## DISCUSSION

Targeting TAMs in tumors is essential, as these cells play a key role in tumor resistance to immune attacks and treatments. Many studies have demonstrated that eliminating TAMs in various mouse tumor models promotes or participates in tumor regression [19–21]. Although mice remain the best model for specific TAMs targeting, developing robust and simple *ex vivo* models to provide an initial pre-clinical proof of concept is a necessity—especially to reduce the number of animals used and to enable high-throughput screening.

We developed a simple and robust 3D model co-culturing lung cancer cells and M2M, which grew for at least seven days recapitulating cancer cell proliferation and maintenance of the M2M population. Moreover, polarized M2 macrophages strongly promoted cancer cell proliferation in this 3D model, whereas monocytes did not, demonstrating that these macrophages exhibit a protumoral function similar to tumor-associated macrophages *in vivo*. This aligns with previous studies showing that monocytes are not fully polarized into M2-like macrophages solely by co-culture with cancer cells [12].

Although many M2M died early in both 3D and 2D co-cultures with cancer cells, we showed that the M2 phenotype of the remaining viable macrophages remained stable over seven days, with a slight increase in the M2 marker CD204. Interestingly, we also detected an increase in both M2 markers, CD206 and FR-β, indicating an exacerbation of the M2 phenotype in 2D co-cultures. Moreover, the viable population of M2M in spheroids was sufficient to promote tumor cell proliferation in both 2D and 3D co-cultures.

We thus demonstrated that adding polarized M2 macrophages to a 3D co-culture is enough to maintain a TAM-like phenotype in these macrophages, accompanied by increased cancer cell proliferation. To confirm the robustness of our 3D model for targeting TAM-like cells, we showed that M1 macrophages co-cultured with A549 cancer cells in 3D for seven days shifted their phenotype toward an M2 phenotype (Figure 3C) and promoted A549 proliferation (Figure 3D). In contrast, M1 macrophages in 2D co-cultures did not show this phenotype shift— except for increased FR-β expression (Supplemental Figure S5)—and this was clearly associated with a decrease in cancer cell numbers, consistent with the well-known antitumor properties of M1 macrophages [22–24].

In this study, we focused on the effective targeting of M2M within a 3D model. We developed a confocal microscopy method combined with spheroid clearing, which allowed us to observe M2 macrophages deep inside the spheroid. Notably, we showed that M2M were distributed throughout the spheroid, including the core. Moreover, we successfully targeted M2M inside the spheroid using specific antibodies such as anti-CD206 and anti-FR-β. Additionally, applying 3D reconstruction approaches to the spheroid images proved to be a valuable tool for obtaining a clear visualization of macrophage status and for more precise cell counting within the spheroid. Finally, using the CSF-1R-targeting drug BLZ945, we specifically induced the death of M2M within this 3D co-culture model.

In conclusion, we have developed and validated a robust 3D lung cancer model that enables the study and targeting of M2M. This model will be especially useful for screening small molecules, antibody-drug conjugates, and other nanotechnologies, such as nanoparticles targeting TAMs, facilitating the selection of promising candidates for further preclinical evaluation in *in vivo* lung cancer models.

## MATERIALS AND METHODS

### Cells

#### Cell lines

A549 and A549-GFP cells were obtained from the Cell Biolabs (USA). These cells were cultured in complete RPMI1640 medium (RPMI 1640 (Gibco, USA), 10% SVF (invitrogen, USA), 100 µg/ml Penicilline-streptomycine (Sigma Aldrich, USA) at 37°C in a humidified 5% CO_2_ atmosphere.

#### Monocyte-derived macrophages

Blood samples from healthy donors were obtained from EFS (Etablissement Français du Sang, Toulouse, France). Peripheral blood mononuclear cells (PBMCs) were collected after separation by Ficoll (Ficoll Paque Plus, GE healthcare, USA) gradient centrifugation. Monocytes were isolated from PBMC by positive selection, using anti-CD14 MicroBeads (Miltenyi, Germany), in accordance with manufacturer’s instructions. To obtain M1 or M2-like macrophages, monocytes were activated respectively with 50 ng/mL of Granulocyte macrophage colony-stimulating factor (GM-CSF, Premium Grade, Miltenyi, Germany) or 25 ng/mL CSF-1 (Premium Grade, Miltenyi, Germany), at day 0 and day 3. At day 6, half of the medium was replaced by fresh medium with 10 ng/mL of interferon γ (IFN-γ, Premium Grade, Miltenyi, Germany) to generate M1 macrophages (M1M), or 25 ng/mL of interleukin-4 (IL-4, Premium Grade, Miltenyi, Germany) and IL-13 (Premium Grade, Miltenyi, Germany) to generate M2 macrophages (M2M). For M1M, 18h after IFN-γ addition, 10 ng/mL of lipopolysaccharide (LPS, Sigma Aldrich, USA) was added in the culture.

### Cultures cells

#### 2D Co-cultures

After polarization, M1M or M2M were detached with Accutase (Bio Legend, USA) at 37°C for 20 min, then labeled with CellTracker Orange (CMTMR, Thermofischer, USA) according to the manufacturer’s instructions. A549 were detached with Trypsin (Gibco, USA) and then labeled with CellTrace Violet (CTV) or CellTrace CFSE (Thermofischer, USA) according to the manufacturer’s instructions. A549 were seeded alone or in co-culture with M1M or M2M at a ratio of 1:1 (0,25 × 10^6^ total cells) in 1.5 mL of complete medium into 12 wells plates and cultured for 7 days at 37°C in 5% CO_2_.

#### Spheroid models

Macrophages were labeled with CMTMR as described in 2D co-cultures. A549 labelled with CellTrace Violet or GFP were seeded alone or in co-culture with M1M or M2M at a ratio of 1:1 (5,000 total cells/well) in a final volume of 200 µL/well in complete medium enriched or not with Matrigel (Corning Matrigel Matrix High Concentration, Sigma Aldrich, USA) into 96-well ultra-low attachment U-bottom plates (Nunclon sphera, Thermofischer, USA). To favour the spheroid formation, plates were centrifuged for 10 min at 215g, then incubated for at least 3 days at 37°C in 5% CO_2_.

### Cancer cell proliferation analysis

At days 3, 5, and 7 after the start of the culture, cells were counted by trypan blue assay on a Malassez counting chamber. For the 3D model, spheroids were pooled and dissociated by Accutase before the counting on a Malassez counting chamber. The cancer cells number was determined by multiplying the percentage of CTV^+^/CMTMR^-^ or CFSE^+^/CMTMR^-^ cells obtained by flow cytometry by the count number.

### Cell viability assays

After M2M polarization, BLZ945 was added at increasing concentrations (0.1, 1 and 10 µg/mL). After 4 days, the percentage of viability was measured with the Cell Meter Cell Viability Assay kit *Green fluorescence* (AAT Bioquest, USA) according to the fabricant protocol. Analysis of the M2M viability inside the spheroids was performed by flow cytometry.

### Microscopy

#### 2D imaging

After 3, 5, 7 days, spheroids were visualized by bright field (BF) and fluorescent microscopy on an automated spinning disk confocal HCS device equipped with a 5x objective (Operetta, Perkin Elmer, France). Morphological parameters (BF area) and macrophage numbers were determined and analyzed with the Columbus software.

#### 3D imaging

Spheroids were incubated with anti-FR-⍰ coupled with CF633 (Mix-n-Stain™ CF™633 Antibody Labeling Kit, Clinisciences, France), anti-CD206 coupled with CF405S (Mix-n-Stain™ CF™405S Antibody Labeling Kit, Clinisciences, France), anti-CD163 coupled with CF405S (Mix-n-Stain™ CF™405S Antibody Labeling Kit, Clinisciences, France), anti-CSF-1 coupled with CF405S (Mix-n-Stain™ CF™405S Antibody Labeling Kit, Clinisciences, France) at 4 µg/mL between 6h to 72h. The spheroids were then fixed directly in the wells with 4% PFA (PFA 16% Solution EM grade, Electron Microscopy Sciences, USA) during 4 h at room temperature. Then the spheroids were cleared with RapiClear 1,49 (SunJin Lab, Nikon, France) for at least 24h between 2 coverslips with two iSpacer of 0.25 mm (Nikon, Japan). Acquisition of images was performed with a LSM880 or LSM980 confocal microscope with 25X objective (Zeiss, Germany).

### Flow cytometry

After detachment and dissociation, cells were resuspended in flow cytometry buffer (2% SVF in PBS) containing 2,5 µg/mL of Human BD Fc Block™ (BD Pharmigen, France) and incubated for 15 min at room temperature. Then, cells were stained with antibodies: anti-CD163-BUV395 (BD Biosciences, USA), anti-CD204-PC7 (Sony Biotechnology, USA), anti-CD206-BV510 (Sony Biotechnology, USA), anti-CD209-PerCP Cy5.5 (BD Biosciences, USA), anti-CD86-BV605 (Sony Biotechnology, USA), anti-CD64-APC Cy7 (BioLegend, USA), anti-FR-β-CF633 (Biotum, USA), at 5 µg/ml, for 20 min at 4°C. Cells were then stained with Annexin-V-FITC according to manufacturer protocol and resuspended in Annexin-V buffer containing 0,8 µg/mL 7-AAD (Sony Biotechnology, USA) to measure the cell viability. All the results were acquired on a Fortessa flow cytometer (BD Biosciences, France) and data were analyzed with Flow Logic 700.2A (Inivai Technologies, Australia).

### 3D spheroid reconstruction from confocal images

Confocal microscopy images were analyzed by IMARIS software (version 10.2, Oxford Instruments). Spots were defined according to a diameter of 8 µm for M2M and 9 µm for antibody labelling. Then the percentage of M2M labelled with anti-CD206, anti-CD163, anti-FR-β was determined according to the distance between spots, with a maximum distance of 6 µm. Finally, the volume was calculated with the surface tool used on the fluorescence of GFP A549.

### 3D Image Analysis in silico

Multichannel 3D fluorescence images of spheroids were acquired in .czi format, each composed of 60 to 93 z-stacks and three fluorescent channels: one for CNMTR^+^ macrophages (red) and two channels of interest (blue and white), corresponding to distinct antibodies (CD163 or CD206, and FR-β, respectively). Image analysis was performed using Fiji (ImageJ v1.54p) with the 3D Objects Counter plugin (v2.0) for object detection and quantification. CNMTR^+^ macrophages, blue, and white objects were segmented independently using the 3D Objects Counter plugin. A minimum object size threshold of 5 voxels was applied to exclude background noise and small artifacts.

Two different analyses approaches were used, differing in pre-processing and thresholding methods:

#### -Automatic thresholding approach

Prior to segmentation, gamma correction was applied to the blue channel to enhance contrast and improve signal clarity. Gamma values were optimized individually for each spheroid. Segmentation was then performed using the default intensity threshold computed automatically by Fiji. This threshold is calculated using the Max Entropy method [25], which determines a global threshold that maximizes the combined entropy of foreground and background pixel distributions across the 3D image histogram.

#### -Empirical thresholding approach

No gamma correction was applied. Instead, segmentation thresholds were manually defined based on visual inspection and adjusted independently for each fluorescence channel to best capture relevant signals.

After segmentation, the 3D centroid coordinates of each object were extracted. To evaluate antibody uptake by macrophages, 3D Euclidean distances were computed between each macrophage centroid and all blue and white object centroids. Macrophage objects located within 6 µm of either a blue or white centroid were considered positive for the corresponding marker. Macrophages located within 6 µm of both blue and white centroids were classified as double-positive.

### Statistical analysis

Results are expressed as the mean ± SEM of at least three independent experiments (one exception in Figure 5). Normality and lognormality tests were performed on the data. Then statistical analysis was performed using one or two-way ANOVA tests or student t-test or Wilcoxon test depending on the normality using Graph Pad Prism 10.1. Differences were considered significant when p < 0.05.

## Author Contributions

CB performed the experiments and analyzed the data. FR performed the in silico experiments. FG, MD and JB helped in the performing of experiments. LL and MF participated in imaging and cytometry experiments. MP, CBo, VP and VG participated in the discussion of the results. CB, VG and MP designed the experiments. MP supervised the study and wrote the paper. All authors have reviewed the submitted version of the manuscript.

## Institutional Review Board Statement

The study was conducted according to the guidelines of the Declaration of Helsinki, and approved by the Institutional Review Board (or Ethics Committee) of INSERM (protocol code DC-2013-1903 and date of approval 2013).

