## Supplemental data for "Targeting human M2 macrophages with antibodies in optimized 3D tumor spheroids"

Supplemental Figure S1

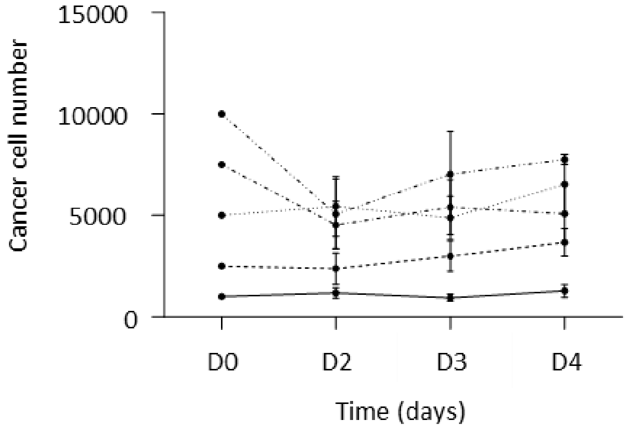


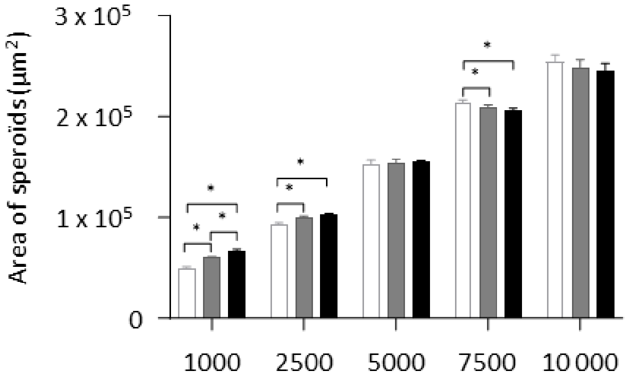


Cancer cell line proliferation in 3D cultures for 4 days at 5 starting points (1000 (n=4), 2500 (n=7), 5000 (n=7), 7500 (n=4), 10 000 (n=4)) and spheroids area (white for day 2, grey for day 3, black for day 4) (n=15)

Supplemental Figure S2

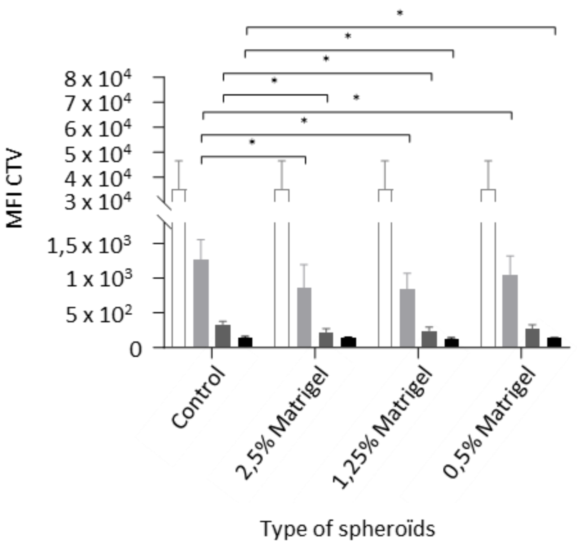


Cancer cell proliferation measured by the CellTrace Violet fluorescence (CTV) dilution at day 0 (white), day 3 (light grey), day 5 (grey) and day 7 (dark grey) of culture.

Supplemental Figure S3

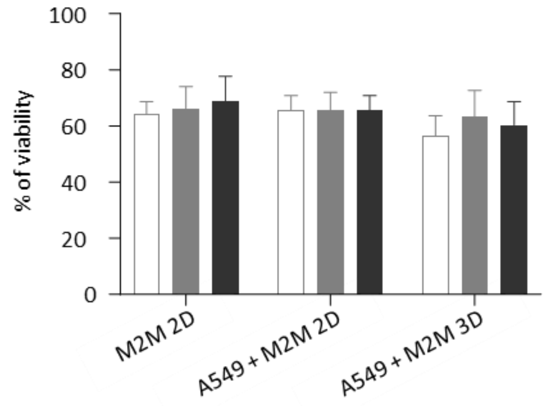


Percent of M2M viability in 2D/3D co-culture with A549 or alone evaluated by Annexin-V/7-AAD labeling in flow cytometry - White day 0, gray day 5, black day 7

Supplemental Figure S4

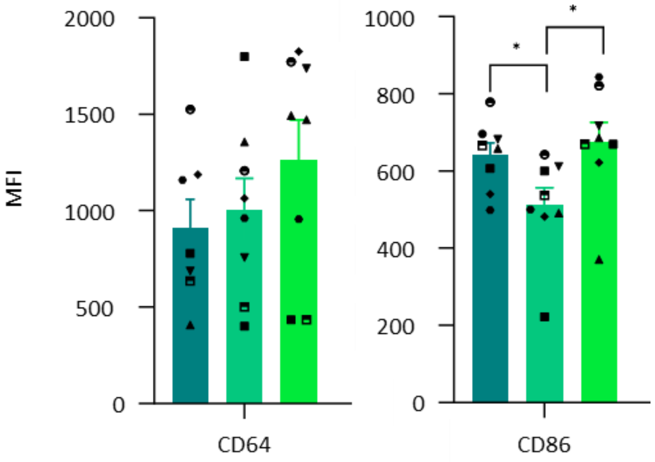


Flow cytometry analysis of CD86 and CD64 expression on M2M at day 7 of culture (8 donors) - dark green M2M alone, green M2M in 2D co-culture with A549, light green M2M in 3D co-culture with A549.

Supplemental Figure S5

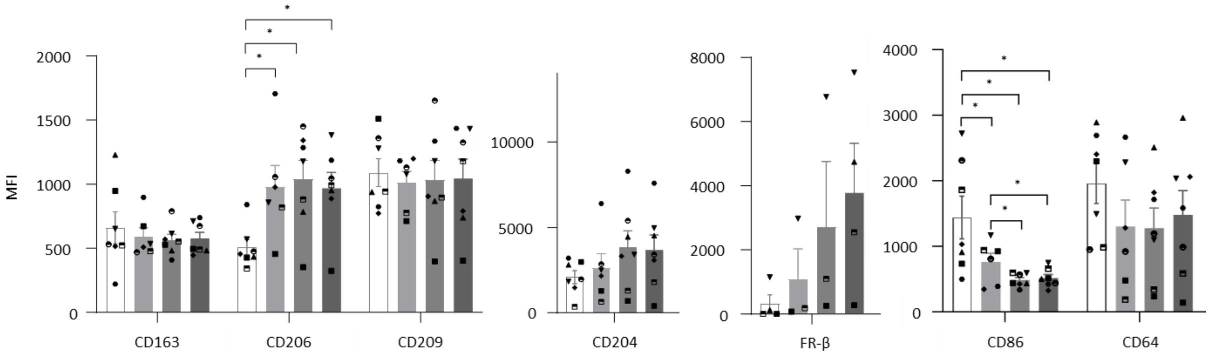


Flow cytometry analysis of markers expression on M1M during 2D cocultures with A549 (6 donors) - White day 0, light grey day 3, grey day 5, black day 7

Supplemental Figure S6

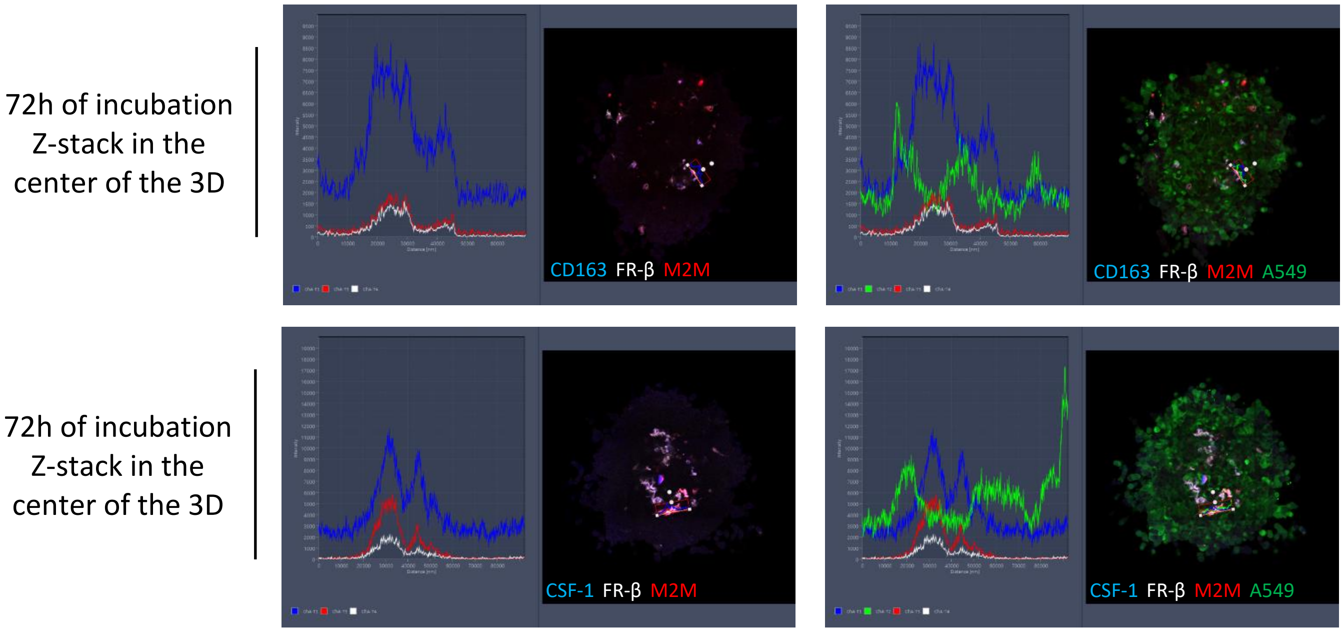


Co-localization profile of the fluorescence of CD163 (blue), CSF-1 (blue), FR-β (white), M2M (red), A549 (green) after 72h of incubation with the different antibodies.

Supplemental Figure S7

IMARIS movie of the modeling of a spheroid with A549 (green) and M2M (red) incubated with anti-CD206 (blue), and anti-FR-β (white) during 3 days
Spots red represents M2M
Spots orange represents the anti-FR-β colocalize with spots of M2M
Spots pink represents the anti-FR-β not colocalize with spots of M2M
Spots blue represents the anti-CD206 colocalize with spots of M2M
Spots green represents the anti-CD206 not colocalize with spots of M2M
Spots white/yellow represents the anti-CD206 + anti-FRB colocalize with spots of M2M
Spots violet represents the anti-CD206 - anti-FRB or anti-FRB - anti-CD206 colocalize with spots of M2M
